# Role of somatostatin-positive cortical interneurons in the generation of sleep slow waves

**DOI:** 10.1101/088443

**Authors:** Chadd M. Funk, Kayla Peelman, Michele Bellesi, William Marshall, Chiara Cirelli, Giulio Tononi

## Abstract

Cortical slow waves – the hallmark of NREM sleep - reflect near-synchronous OFF periods in cortical neurons. However, the mechanisms triggering such OFF periods are unclear, as there is little evidence for somatic inhibition. We studied cortical inhibitory interneurons that express somatostatin (SOM), because ∼70% of them are Martinotti cells that target diffusely layer 1 and can block excitatory transmission presynaptically, at glutamatergic terminals, and postsynaptically, at apical dendrites, without inhibiting the soma. In freely moving mice, we show that SOM+ cells can fire immediately before slow waves and their optogenetic stimulation triggers neuronal OFF periods during sleep. Next, we show that chemogenetic activation of SOM+ cells increases slow wave activity (SWA), the slope of individual slow waves, and the duration of NREM sleep; whereas their chemogenetic inhibition decreases SWA and slow wave incidence without changing time spent asleep. By contrast, activation of parvalbumin+ (PV+) cells, the most numerous population of cortical inhibitory neurons, greatly decreases SWA and cortical firing. These results indicate that SOM+ cells, but not PV+ cells, are involved in the generation of sleep slow waves. Whether Martinotti cells are solely responsible for this effect, or are complemented by other classes of inhibitory neurons, remains to be investigated.

## RESULTS AND DISCUSSION

Non-rapid eye movement (NREM) sleep is characterized by spindles and slow waves. While the mechanisms responsible for the generation of spindles are well characterized [1], those underlying slow waves remain unclear [2]. Slow waves are generated in the cerebral cortex even when isolated from thalamic input [1], although the thalamus is important for their full expression [3]. Every second or so in the course of NREM sleep, when the EEG records the occurrence of slow waves, cortical cells undergo a sharp hyperpolarization of the membrane potential lasting for tens to hundreds of msec (down state) during which neurons are silent (OFF period). This is followed by a return to a tonic depolarization (up state) often accompanied by neuronal firing (ON period). The near-synchronous occurrence of down states in large sectors of the cortex is detected in the EEG as the negative peak that typifies each slow wave. It was suggested that the OFF periods that characterize NREM sleep may occur, upon a background of increased potassium leak currents, through a passive “disfacilitation” associated with reduced synaptic input [1]. However, it is not clear what would trigger a decrease in synaptic input in the first place. Moreover, it is difficult to explain why the occurrence of the OFF periods is remarkably sharp and synchronous across large populations of neurons [4]. Finally, if reduced activity were the primary trigger of slow waves, it would be hard to explain why they can be induced reliably by electrical [5] or transcranial magnetic stimulation [6]. An alternative possibility is that down states may be induced by active inhibition [7, 8]. However, except for a small fraction of pyramidal neurons [7], intracellular recordings in the cell body of pyramidal cells are consistent with disfacilitation rather than with direct inhibition [1], and there is currently no causal evidence for the involvement of specific inhibitory cell types in the generation of slow waves.

We reasoned that a prominent class of cortical inhibitory interneurons that express somatostatin (SOM) – Martinotti cells – have several features that could explain the synchronous induction of OFF periods and associated slow waves in the absence of somatic inhibition. First, they are found throughout the cortex, where they target especially layer 1 (L1), the site of termination of many cortico-cortical connections, particularly back-connections, and of diffusely projecting thalamocortical projections from matrix cells [9, 10]. In L1, SOM+ cells powerfully inhibit excitatory transmission among pyramidal cells [11] through synaptic spillover over presynaptic GABAb receptors on glutamatergic terminals, which could explain the profound disfacilitation at the cell body despite little evidence for inhibitory currents [1]. Moreover, in L1 SOM+ cells inhibit distal dendrites of pyramidal cells [12] through GABAa [13] and likely GABAb receptors [14, 15]. Martinotti cells have high density of connections and convergence onto pyramidal cells, irrespective of specific subnetworks, and can thus provide indiscriminate pyramidal inhibition [16, 17]. Furthermore, SOM+ cells are activated by strong synchronous firing of pyramidal cells [18, 19] through progressively facilitating synapses [12, 20]. When many SOM+ cells are activated together, they act as ‘master regulators,’ inhibiting all other cell types but themselves [21], in line with the observations that OFF periods are found across cell types [1]. SOM+ cells tend to fire more strongly late during the up state, and have been implicated in the termination of the up state in computer simulations [22] and in vitro [20], possibly by activating GABAb receptors [14, 23]. Finally, Martinotti cells have uniquely broad and complex axonal arborizations [23], which could account for the broad synchrony of sleep slow waves [1], and they form an electrical syncytium through gap junctions [19, 24], which could favor the spread of slow waves in cortex and possibly account for their traveling nature [25].

There is currently no molecular marker to target Martinotti cells exclusively [26]. Instead, we used SOM-Cre mice [27], because Martinotti cells form the bulk (∼70%) of SOM-expressing cells in cortex [21]. Adeno-associated virus (AAV)-driven expression of Cre-dependent enhanced green fluorescent protein (eGFP) in SOM-Cre mice confirmed that SOM+ cells have strong projections to L1 both locally and up to several millimeters away from the injection site (Suppl. Figure S1A), consistent with the known projections of Martinotti interneurons [23]. We then injected SOM-Cre mice either in one frontal site (secondary motor cortex, M2) with Cre-inducible AAV expressing channelrhodopsin-2 (ChR2), or bilaterally in frontal (M2) and parietal cortex with Cre-inducible AAV expressing the designer receptors hM3Dq or hM4Di. After ∼3 weeks to allow for viral expression, mice were implanted with frontal and parietal EEG electrodes and intracortical laminar probes in M2, and continuously recorded for several days across multiple sleep-wake cycles, during sleep deprivation and subsequent recovery sleep, as well as during and after optogenetic or chemogenetic stimulation (Suppl. Figure S1B).

First, we characterized the firing pattern of 3 SOM+ cells in M2 that were identified by optogenetic tagging in freely moving animals (Figure 1A) and whose activity could be followed for several days during the sleep-wake cycle (Figure 1B). All 3 cells exhibited state-specific firing modulation, with higher firing levels during active wake with exploration relative to quiet wake (consistent with [28], but see [29]) and during NREM sleep relative to REM sleep (Figure 1C). Two of these cells were also recorded during recovery sleep after sleep deprivation, when they fired more than during baseline sleep (Figure 1D). In these cells firing was negatively correlated with spindle activity (Suppl. Figure S2A) and positively correlated with SWA, both in baseline and after sleep deprivation (Figure 1E). In all 3 cells firing significantly increased between 50 and 100 ms before the onset of locally detected slow waves, consistent with a link between SOM+ cell firing and SWA in natural sleep (Figure 1F). Although correlative and restricted to a few neurons, these results are consistent with the hypothesis that SOM+ cell firing may be linked to the occurrence of slow waves in natural sleep.

**Figure 1.**
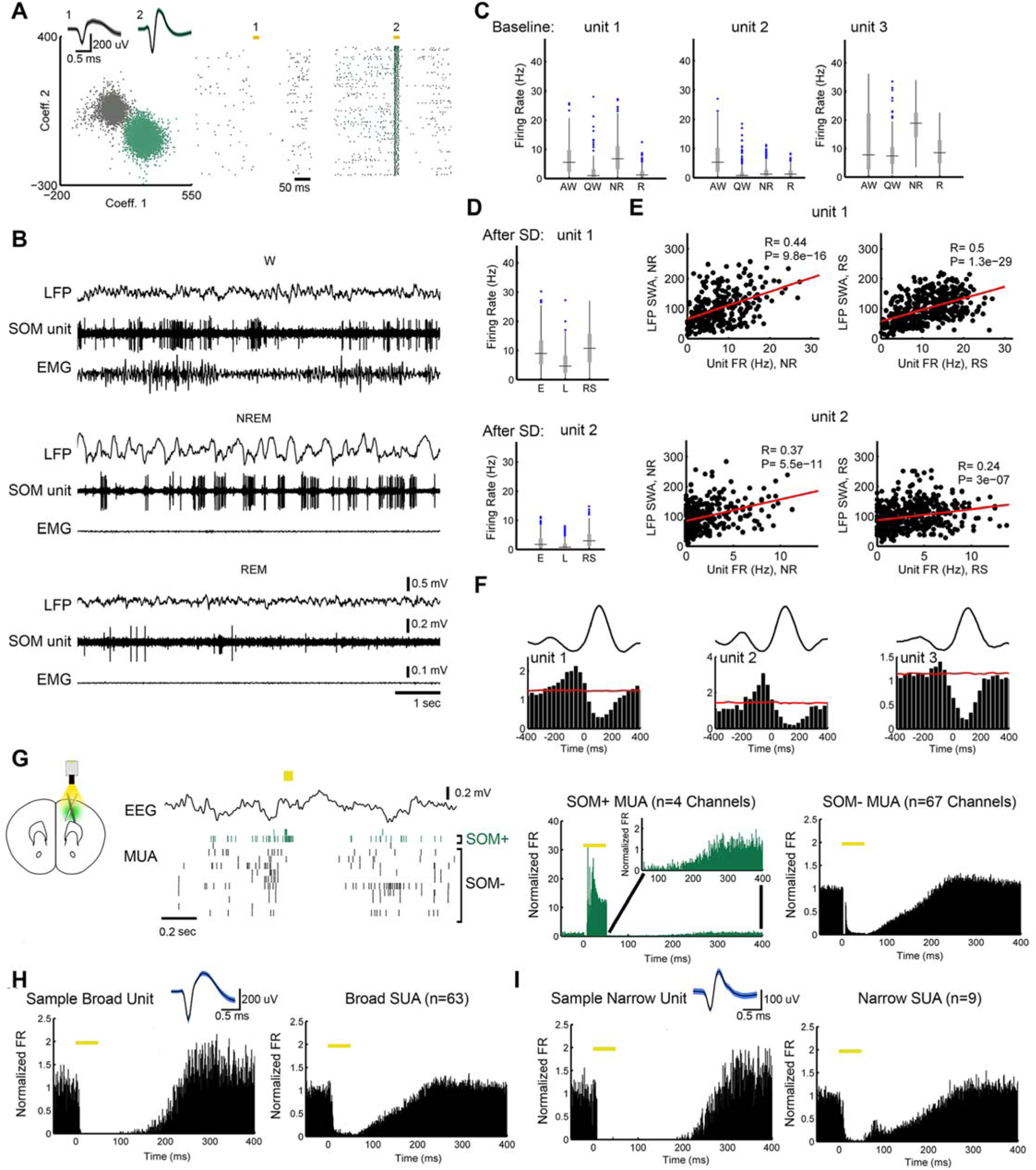
Firing of SOM+ cells is locked to slow waves and triggers OFF periods during sleep. **A**, Clustering results for one well-isolated unit, in green, tagged by ChR2 stimulation (putative SOM+). Yellow bars indicate the time of laser stimulation (20 msec). **B,** Raw LFP and activity from the same unit; **C**, Mean firing rates (light period) of the 3 tagged units; AW, QW, active, quiet wake. **D**, Mean firing rates in recovery sleep (RS) after sleep deprivation compared to early and late NREM sleep during baseline (E,L). **E**, Correlation between unit firing and SWA in baseline sleep (NR) and recovery sleep after SD (RS). **F,** Firing is locked to local slow waves and increases significantly (above red line) before their onset (0 msec, defined by the zero crossing of each slow wave, see Methods; firing normalized for mean in NREM, red line is 99^th^ percentile FR for simulated spike trains with shuffled ISIs). **G**, Left, example of MUA changes in one responding (SOM+, green) channel and several SOM-channels after laser stimulation (yellow bar, 50 msec); right, mean MUA changes for all SOM+ and SOM-channels (7 mice). Inset, expanded Y axis to show the firing of SOM+ channels after laser pulse**. H,** Left, change in firing in one broad unit (putative excitatory, spike waveform shown on top); right, mean of all broad units. **I**, as in **H** for narrow (putative inhibitory) units.

To directly test whether SOM+ cell firing is sufficient to induce OFF periods in the local cortical network of freely moving animals, we used optogenetic stimulation to briefly activate SOM+ cells in M2 during natural sleep, while recording multi-unit activity (MUA) and single unit activity from the same area. SOM+ cell activation with laser pulses (50 msec) consistently induced OFF periods in MUA that outlasted the duration of the pulse and were comparable in length to those occurring during stimulation-free sleep (190 ± 32 ms vs. 174 ms ± 10 ms; mean ± sd). OFF periods occurred in all recorded M2 channels, including a majority whose MUA decreased during the pulse and remained decreased in the subsequent OFF period, and a few channels whose MUA sharply increased during the pulse. These “responding” channels, which presumably contained at least one SOM+ cell, accounted for ∼ 7% of all channels, consistent with the number of cortical cells that are SOM+ [30] (SOM+ channel, Figure 1G). After spike sorting 72 single units were identified across all channels in superficial and deep layers, including 63 broad (putative excitatory) units, and 9 narrow (putative inhibitory) fast-spiking neurons. All units were effectively silenced by SOM+ cell activation, in line with the ability of SOM+ cells to inhibit all other cortical cell types [21] (Figure 1H,I). Diffuse and persistent silencing was also obtained with shorter (20 msec) laser pulses (data not shown). Thus, multiunit OFF periods induced by SOM+ cell activation appear similar to those of physiological NREM sleep and crucially, brief optogenetic activation of SOM+ cells silences all recorded single units for a period that outlasts the stimulation – a characteristic feature of the bistability of slow oscillations [31].

Next, we tested whether selective chemogenetic activation of SOM+ interneurons over a large portion of cortex (bilateral frontal + parietal) would promote a prolonged increase in SWA. We chose a chemogenetic approach because it allows for sustained depolarization of SOM+ cells without enforcing highly synchronous activation as in optogenetic experiments. To control for its possible sedative effects, the selective ligand clozapine-N-oxide (CNO) was injected in SOM+ mice expressing the excitatory receptor hM3Dq AAV, and in other mice expressing non-hM3Dq AAV (controls). The effects of SOM+ cell chemogenetic activation on behavioral state and spectral activity in specific frequency bands were assessed using linear mixed effect (LME) models that included mouse as a random effect, as well as day (baseline, CNO), condition (group, e.g. hM3Dq, non-hM3Dq), and time (hour) as categorical fixed effects. Specifically, we tested for a significant condition by day interaction, since we hypothesized that there would be differences between groups on the CNO day but not on the baseline day. If a significant interaction was detected, post-hoc tests (adjusted for multiple comparisons) were used to confirm whether a significant increase or decrease was only present during the CNO day. With this approach, we could isolate the specific effects of SOM+ cell activation above and beyond the effects due to CNO injection alone (controls). The same statistical approach was used for SOM+ cell inhibition and PV+ cell activation experiments (see below).

First, we confirmed that CNO administration in SOM-Cre mice expressing hM3Dq drove activation of Fos -a marker of neuronal activity - in SOM+, hM3Dq+ neurons, indicating that the drug was able to activate its target cells in vivo (Figure 2A). Next, SOM-Cre mice previously injected with either hM3Dq AAV or non-hM3Dq AAV (n=6 mice/group) were administered CNO (5mg/kg) midway through the light phase, when most sleep pressure has dissipated. After CNO injection mice continued to cycle through sleep-wake states and NREM sleep was characterized by decreased muscle tone, slow waves in both frontal EEG and LFP signals, and OFF periods in MUA (Figure 2B,C), suggesting that qualitatively, physiological sleep continued following SOM+ cell activation. Quantitatively, relative to controls, SOM+ cell activation resulted in a sustained increase in NREM SWA in both the EEG (p < 3e-5) and M2 LFP (p = 3.6e-10) that lasted for the rest of the light cycle (Figure 2D). Further spectral analysis in M2 found a broad shift toward lower frequency activity during NREM sleep, with a significant increase in SWA, theta (6-8 Hz), and alpha (8-15 Hz) bands and a decrease in beta (15-30 Hz), low (30-60 Hz) and high (60-100 Hz) gamma bands (beta, p = 0.028; alpha, p = 0.002; all others p < 1e-8 for Hrs 7-12; Figure 2E). There was no difference in spindle power between groups (p=0.93; Suppl. Figure 2B) and a significant decrease in spindle incidence (p = 2.9e-6; Suppl. Figure 2C). Similar effects – increased SWA, alpha, theta and beta power and decreased low and high gamma power – were present in REM sleep (p < 1e-7; Figure 2E). In wake, SWA increased (p < 0.0005) and high gamma decreased (p = 6.9e-6; Figure 2E). Sleep architecture was also modulated by SOM+ cell activation, with a trend towards increased NREM sleep and a significant decrease in wake and REM sleep (Figure 2D).

**Figure 2.**
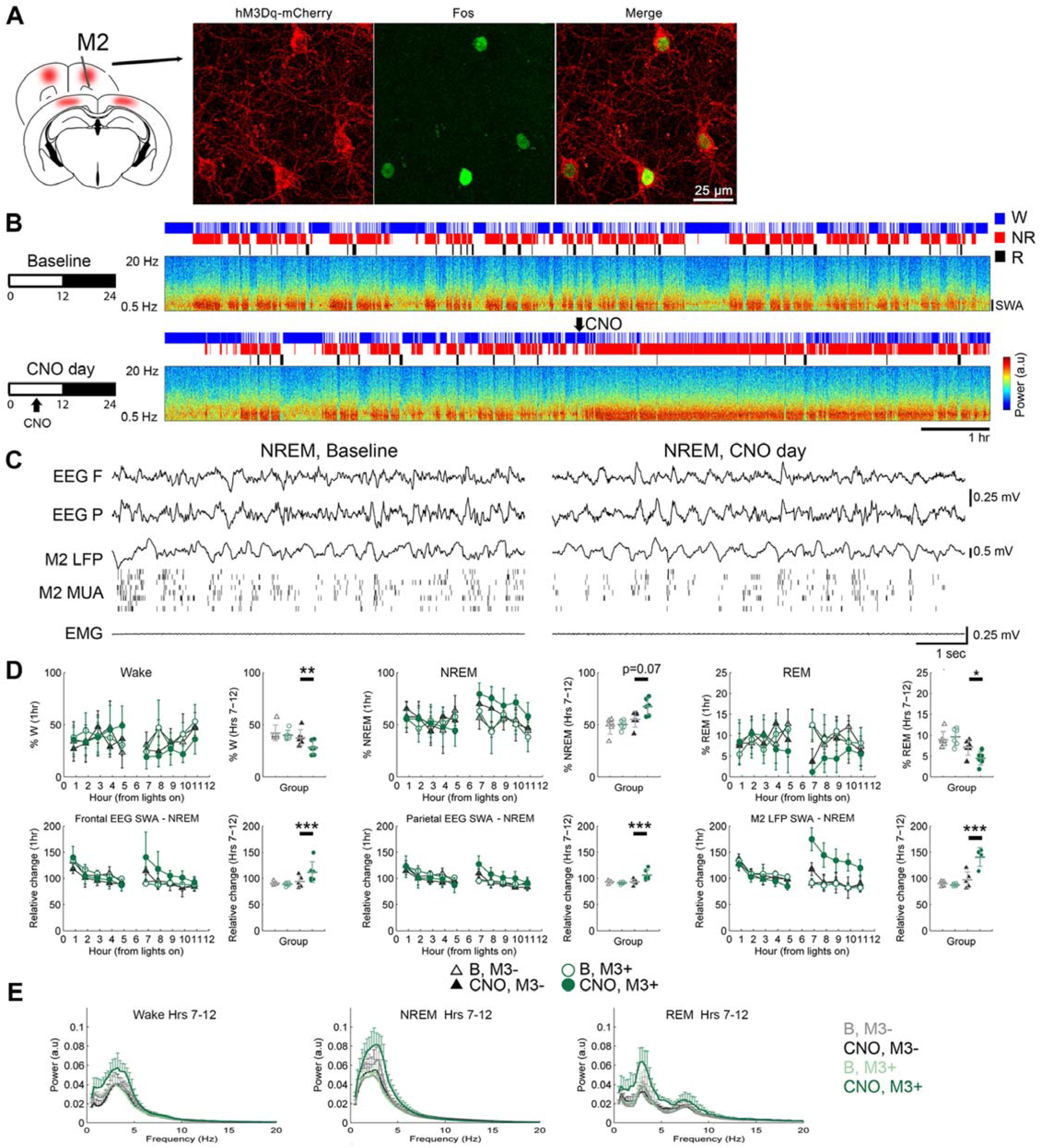
Chemogenetic SOM+ cell activation leads to a sustained increase in SWA. **A**, Fos induction in SOM+, hM3Dq+ neurons after CNO. **B,** Effects on sleep and SWA (NREM sleep EEG power, 0.5-4.5 Hz, M2) in one representative SOM-Cre hM3Dq+ mouse given CNO (5 mg/kg i.p) in the middle of the light phase. Hypnograms and spectra shown for entire light period (12hrs). **C**, Raw traces from the same animal on baseline and post-CNO. **D,** Sleep/wake and SWA percentages in baseline (B) and after CNO in mice expressing hM3Dq AAV (M3+, 6 mice) and mice expressing non-hM3Dq AAV (controls, M3-, 6 mice). For each parameter the left panel shows the hour-by-hour data, which were used to run the LME models, locked to first sleep bout after the start of the light period and after midpoint of the light period (when CNO was given); if a significant condition (group) by day interaction was found, post-hoc tests were run to isolate the main effect of CNO (right panel). *, p<0.05; **, p<0.01; ***, p<0.001. **E,** Power spectra for the second half of the light period in each vigilance state.

To compare the effects of SOM+ cell activation to well-known procedures that lead to increased sleep pressure and SWA, we subjected the same SOM-Cre/AAV-hM3Dq animals to sleep deprivation through exploration of novel objects for 6 hours during the first half of the light phase, a period during which mice are normally asleep. Consistent with previous studies [32], we found that sleep deprivation led to an early (first hour of recovery) increase in SWA (p = 2e-16), theta (p = 2e-16), alpha (p = 2.2e-15) and beta activity (p = 0.007) during NREM sleep and in theta activity during wake (p = 2.8e-6), as well as to an increase in time spent in NREM sleep (Suppl. Figure 3A,B). Thus, the enhancement of slow frequencies after SOM+ cell activation resembles that induced by sleep deprivation. The incidence and slope of the slow waves – two additional, highly sensitive measures of sleep pressure [5] – also increased in M2 after both sleep deprivation and SOM+ cell activation (Suppl. Figure 3C-E). Moreover, analysis of laminar recordings in baseline, recovery NREM sleep, and NREM sleep following chemogenetic activation showed that in all cases slow waves were largest in the LFP recordings from deep layers and were accompanied by OFF periods across all layers (Suppl. Figure 3F), consistent with previous reports from NREM sleep [33]. Thus, like sleep deprivation, chemogenetic activation of SOM+ cells promotes the occurrence of slow waves that electrophysiologically resemble those in natural NREM sleep and also increases their slope, a sign of enhanced neuronal synchrony [5]. Altogether, these results suggest that the stimulation of SOM+ cells may promote the same physiological mechanisms underlying sleep slow waves.

**Figure 3.**
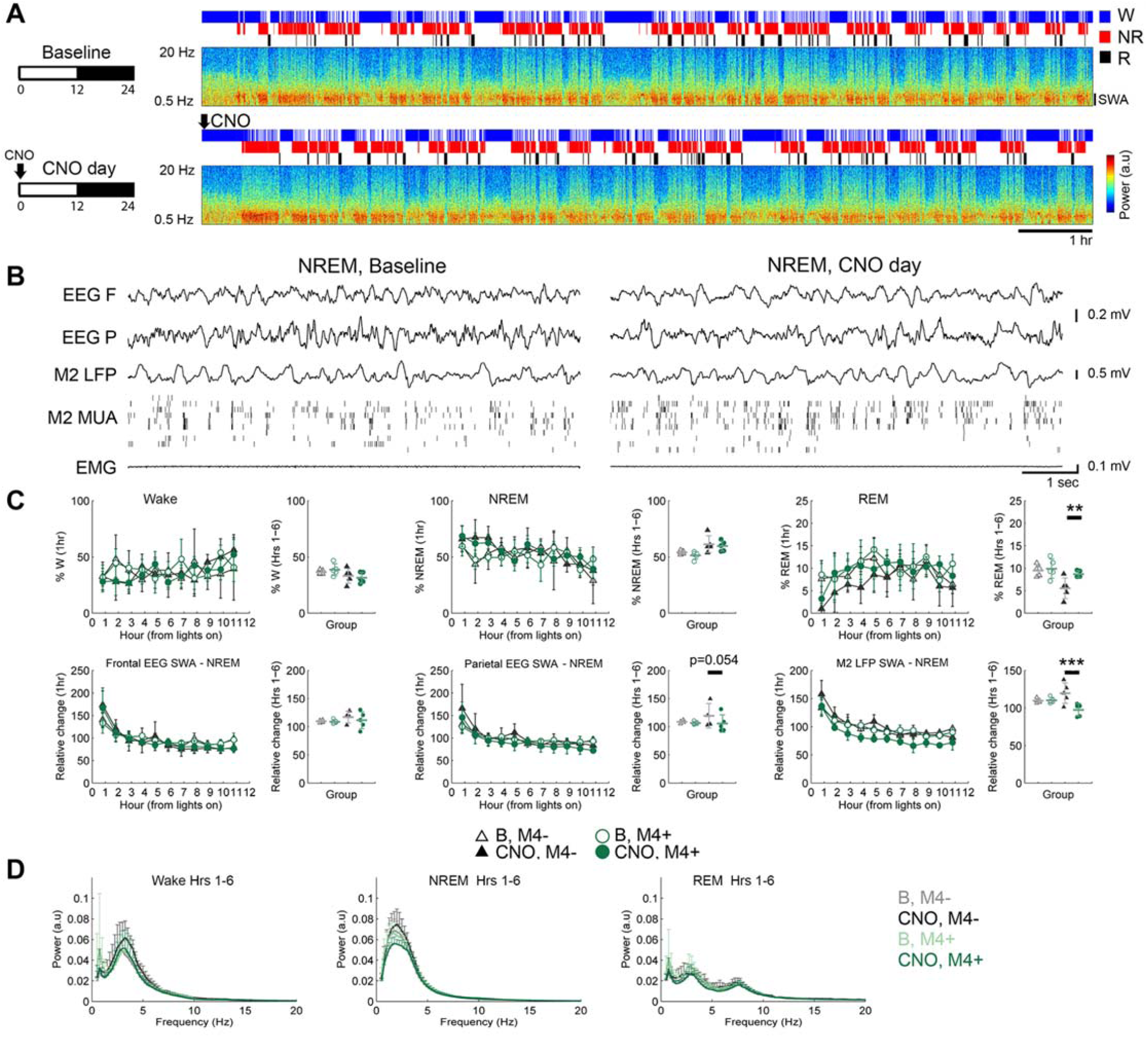
Chemogenetic inactivation of SOM+ cells leads to a sustained decrease in SWA. **A-B**, Effects on sleep and SWA in M2 during the light period in one representative SOM-Cre hM4Di+ mouse given CNO (10 mg/kg i.p) at light onset, and raw traces from the same animal in the second hour post-injection. **C,** Effects on SWA and behavioral states after CNO (M4+, 5 mice; controls, M4-5 mice). For each parameter the left panel shows hour-by-hour data locked to the first sleep bout after the start of the light period (when CNO was given), while the right panel shows the mean values from the first half of the light period. LME models were run on hour-by-hour data; if a significant condition (group) by day interaction was found, post-hoc tests were run to isolate the main effect of CNO (right panel). **, p<0.01; ***, p<0.001. **D,** Power spectra for the first half of the light period in each vigilance state.

Other SOM-Cre mice were injected with the inhibitory receptor hM4Di (M4+, n=5 mice) or non-hM4Di AAV (M4-, controls, n=5 mice). After CNO, hM4Di+ mice did not differ from controls in the time spent awake or asleep. By contrast they showed, during NREM sleep, a persistent decrease in SWA (Figure 3A-D), associated with a decrease in the incidence of slow waves (N/min, 33.4 ± 2.0 vs. 29.9 ± 2.7, p = 0.01, mean ± sd) and during REM sleep, a broad decrease in power across most frequencies (Figure 3D).

Previous work has described a group of cortical neurons containing high levels of neuronal nitric oxide synthase - type I nNOS+ cells – that expresses Fos during early recovery sleep, when SWA is high [34]. We first confirmed this finding in 2 mice, in which we found that after 90 min of recovery sleep following 6 hours of sleep deprivation most nNOS+ cells were also Fos+ (61%), and were mainly located in deep layers, as previously reported [35] (data not shown). Since all type I nNOS+ cells are also SOM+ cells but not vice versa, [35] in other mice (n=6) we assessed Fos expression in SOM+ cells and found that only a small subset of SOM+ neurons were Fos+ during recovery sleep. Conversely, many Fos+ cells during recovery sleep were not SOM+ cells, and Fos+ cells were found in all cortical layers (Suppl. Figure 4). It should be noted that the mean firing rates of most cortical cells are only slightly decreased throughout sleep relative to wake and the relationship between spontaneous neuronal activity and Fos expression is not straightforward [36].

**Figure 4.**
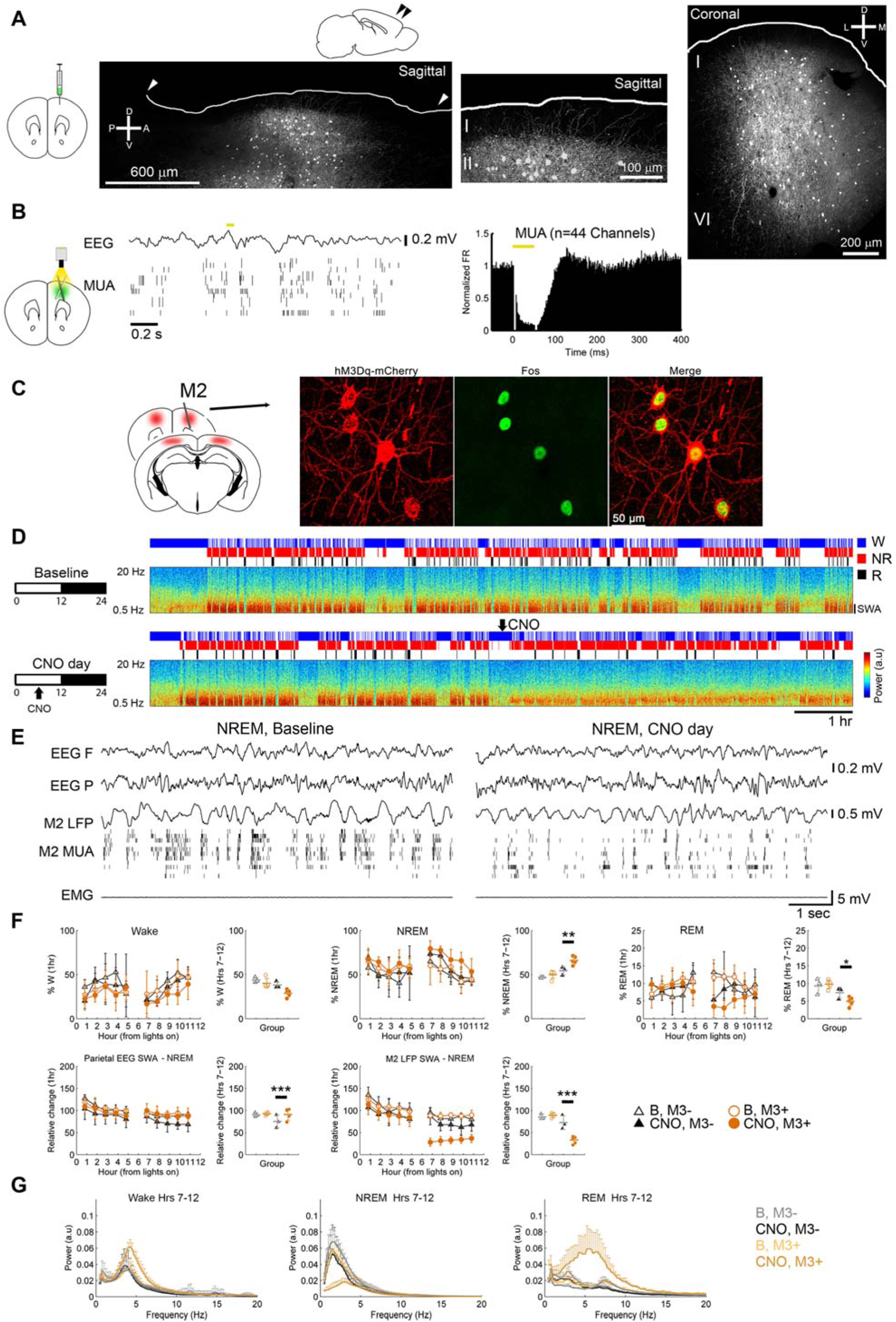
PV activation decreases SWA. **A**, Cortical projections shown in sagittal and coronal sections from a representative PV-Cre mouse injected with AAV1.CAG.Flex.eGFP.WPRE.bGH, with sparse innervation in layer 1. **B**, Optogenetic stimulation of PV+ cells leads to transient suppression of MUA (5 mice). **C**, Fos induction after CNO. **D,E,** Increase in NREM sleep percentage but decrease in M2 SWA following CNO in one representative mouse given CNO (5 mg/kg i.p) in the middle of the day, and raw traces from the same animal during the second hour post-injection. B, baseline. **F,** Effects on SWA and behavioral states (M3+, 5 mice, controls, M3-3 mice). For each parameter the left panel shows hour-by-hour data, which were used to run the LME models, locked to the first sleep bout after the start of the light period and after midpoint of the light period (when CNO was given); if a significant condition (group) by day interaction was found, post-hoc tests were run to isolate the main effect of CNO (right panel). *, P<0.05; **, p<0.01; ***, p<0.001. **G**, Power spectra for the second half of the light period in each vigilance state.

Is the induction of physiological slow waves and NREM sleep specific to SOM+ neurons? The most numerous population of cortical inhibitory cells are parvalbumin (PV) positive [30]. Using a PV-Cre line, we first confirmed that unlike SOM+ cells, PV+ neurons have local projections and their axons mostly avoid L1 (Figure 4A). Optogenetic stimulation of PV+ cells produced a more transient silencing in response to laser pulses than did optogenetic activation of SOM+ cells (Figure 4B; latency to first spike post-pulse, mean ± sd; SOM+ 110 ± 38 ms; PV+ 50 ± 15 ms, p=0.007; 7 SOM+ mice and 5 PV+ mice). Next, we confirmed that CNO induces Fos expression in PV+, hM3Dq+ neurons (Figure 4C), as it does in SOM+, hM3Dq+ cells. Relative to controls injected with non-hM3Dq+ AAV, chemogenetic activation of PV+ cells led to a marked decrease in M2 SWA during NREM sleep (Figure 4D-F, n=5 mice) as well as in all other major frequency bands (all p<0.05; Figure 4G). PV+ cell activation also increased the duration of behaviorally and polygraphically defined NREM sleep and decreased the duration of REM sleep (Figure 4F). Consistent with this broad power decline, during chemogenetic activation of PV+ cells cortical firing was profoundly suppressed during NREM sleep (54 ± 8% decline in MUA, hours 8-12 post-CNO vs. hours 1-6 pre-CNO), much more than after activation of SOM+ cells (27 ± 6% decline; p=0.0009). In contrast, PV+ cell activation led to a broad increase in EEG power during REM sleep (from SWA to beta, all p<0.05; Figure 4G) and during wake (from SWA to alpha, all p<0.05; Figure 4G).

These findings show that slow waves – the hallmark of NREM sleep – are positively regulated by a population of SOM+ cortical inhibitory interneurons. SOM+ cells fire just before the occurrence of slow waves and their acute optogenetic activation produces a neuronal OFF period across putative excitatory and inhibitory neurons, consistent with the broad connectivity of SOM+ cells. Sustained chemogenetic activation of SOM+ cells enhances SWA and the slope of individual slow waves, while their chemogenetic inhibition decreases SWA and slow wave incidence. In contrast, PV+ cell activation profoundly suppresses SWA and cortical activity in general, demonstrating that the positive role SOM+ cells in the generation of sleep slow waves is not shared by all GABAergic cortical interneurons.

Altogether, SOM+ cell activation increased sleep SWA, SOM+ inhibition decreased SWA, and PV+ cell activation nearly suppressed SWA. However, only PV+ cell activation led to a significant increase in NREM sleep duration. Previous studies [32, 37, 38] have also shown that changes in the “intensity” of NREM sleep, as reflected by SWA, can be dissociated from changes in its duration. For instance, several drugs and different regimes of sleep deprivation can affect either NREM sleep or slow waves, or change both but in opposite directions [32, 37]. Moreover, slow waves can change globally or locally in response to wake-induced changes in cortical activity and plasticity, often without any effect on NREM sleep duration [32, 37, 38].

It is possible that in addition to SOM+ cells, other inhibitory cell types play a role in the induction of slow waves. For example, neurogliaform cells can induce indiscriminate GABAb-mediated hyperpolarization through volume transmission [30, 39], although their effects on synaptic strength are less powerful than those of SOM+ cells [11]. The widespread L1 projections of Martinotti cells, which comprise ∼70% of SOM+ cells, makes them a strong candidate for mediating the induction of sleep slow waves reported here, but definitive evidence will require repeating these experiments in Martinotti-specific Cre lines. Future experiments should establish the contribution of other cortical subpopulations of SOM+ cells [40], such as non-Martinotti cells in L4 [41], and other SOM+ cells outside the cortex, including the bistratified SOM+ cells in the hippocampus [42]. Moreover, SOM+ cells in the basal forebrain inhibit all types of neighboring wake-promoting neurons and their optogenetic activation slightly increases NREM sleep, consistent with the finding that a minority of these cells is more active during NREM sleep [43]. Another question is how the activity and excitability of SOM+ cells is controlled by subcortical wake and sleep promoting systems, either directly or through other cortical cell types. SOM+ cells are inhibited by cortical VIP interneurons [44] that fire during locomotion and are themselves under the control of arousal systems [45] and by PV+ interneurons in the basal forebrain, which are wake-on and promote wake [43]. Also, noradrenaline, a neuromodulator that is released at high levels in wake and low levels in sleep, reduces the effectiveness of synapses between SOM+ and pyramidal cells [46] and the conductivity of gap junctions [47], possibly preventing the induction and synchronization of sleep slow waves. Finally, an open question is whether the role of SOM+ cells in promoting SWA may be related to their suggested role in the maturation of cortical circuits and in synaptic plasticity [48].

## Acknowledgements

Funded by NIMH grant R01MH099231 to CC and GT, NINDS grant P01NS083514 to CC and GT, R01GM116916 to GT, Wisconsin Distinguished Rath Graduate Fellowship to CMF, NIGMS T32 GM008962 to CMF. CNO was obtained from NINDS as part of the Rapid Access to Investigative Drug Program.

